# Chemical Characterization of Interacting Genes in Few Subnetworks of Alzheimer’s Disease

**DOI:** 10.1101/364802

**Authors:** Antara Sengupta, Hazel Nicolette Manners, Pabitra Pal Choudhury, Swarup Roy

## Abstract

A number of genes have been identified as a key player in Alzheimer’s disease (AD). Topological analysis of co-expression network reveals that key genes are mostly central or hub genes. The association between a hub gene and its neighbour genes can be derived easily using relative abundance of their expression levels. However, it is still unexplored fact that whether any hub and its neighbour genes within a sub-network exhibits any kind of proximity with respect to their chemical properties of the DNA sequences or not, that code for a sequence of amino acids.

In this work, we try to make a quantitative investigation of the underlying biological facts in DNA sequential and primary protein level in mathematical paradigm. It may gives a holistic view of the interrelationships existing between hub genes and neighbour genes in few selective AD subnetworks. We define a mapping model from physicochemical properties of DNA sequence to chemical characterization of amino acid sequences. We use distribution of chemical groups present in a sequence after decoding into corresponding amino acids to investigate the fact that whether any hub genes are associated closely with its neighbour genes chemically in the subnetworks. Interestingly, our preliminary results confirm the fact the dependent genes that are coexpressed with its hub gene are also having proximity with respect to their amino acid chemical group distributions.

**CCS Concepts:** •Applied computing → Computational genomics;

## 1. INTRODUCTION

Alzheimer’s disease is a neuro-degenerative disease. AD is a type of dementia that causes problems with memory, thinking and behavior. Identifying any key genes and its underlying interaction network may put light on the disease mechanism. In turn it may help design effective therapeutic drug molecules for AD. It has been observed that such key genes or possible regulators are central genes [2], also called hub genes. Studies [1] reveals few genes namely, Amyloid Precursor Protein (APP), PreSenilin1 (PSEN1) and PreSenilin2 (PS2), Apolipoprotein E (ApoE) are some of the key genes in AD. Instead of single gene, generally a set of interacting genes forming a sub-network or module [17, 16] act towards disease abnormalities inside cell. The common way to identify such key genes and its subnetworks is to infer network computationally from microarray gene expression data [15, 3]. Drawing a relationship among pairs of genes in a co-expression network based on certain correlation measures is not always a conclusive step. We feel that similarity based on chemical properties between interacting genes also plays important role during interaction. We try to investigate the chemical proximity of any hub gene and its immediate neighbours within the subnetwork of the hub gene. As a case study, we consider AD responsible few key genes based on their centrality in the co-expression network. We first quantify DNA sequence of a hub gene and its neighbours with respect to their physicochemical properties. Finally, we compute proximity of a hub gene with its neighbours to show that hub genes are closely associated with their neighbours bio-chemically.

We organize the rest of the paper as follows. At first in Section 2 we describe the process of extracting hub genes and sub-networks. In Section 3, we develop a mathematical model to map and arbitrary DNA sequence to its corresponding chemical properties. We compute the proximity of each neighbour genes with all candidate hub genes to see how they are associated with each other (Section 4). Few experimental results are reported in Section 5. Finally, we summarize our work in Section 6 with concluding remarks.

## 2. EXTRACTION OF ALZHEIMER’S SUB-NETWORKS

To perform our experiment we use the gene expression data of Alzheimer’s disease (GSE1297) from the NCBI^1^ data repository. In this datatset, 31 microarrays are used to analyse 9 control and 22 AD subjects of differing severity in the disease and test their correlation. In order to extract sub-networks from microarray gene expression data, we first perform network construction. Any two genes that have similar expression patterns will have lesser distance (higher similarity) and are more likely to interact with each other. Soft thresholding is then applied where interactions having a distance score above a certain threshold are removed. We use a simple parametric distance measure to compute the proximity between two gene expressions.

### Definition 2.1 (PROXIMITY)

*Given two expression vectors x* =*< x*_1_*,x*_2_, ⋯ *x_n_ > and y* =*< y*_1_*,y*_2_, ⋯ *y_n_ > for the genes x and y, the distance, δ*(*x, y*)*, between two genes can be calculated by taking the normalized difference of standard deviations between the two expression profiles*.

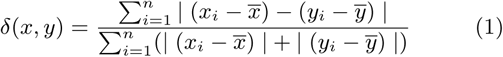

After the network construction, we then extract sub-networks using a density-based clustering approach [14]. These sub-networks are then analysed biologically as well as topologically. To determine how much each of the sub-networks may contribute to Alzheimer’s disease we validate using KEGG pathway [8] analysis. We select sub-networks where the percentage of genes that participate in the AD pathway are considerably high along with low p-value. From these selected sub-networks, identification of hub genes is then done based on Maximum Clique Centrality(MCC) score[4]. MCC score considers the degree of connectivity of a node as well as the size of the branches that it connects to the rest of the network. Genes having high ranking MCC scores in the sub-networks are considered hub genes. MCC score of a node *v* is defined as follows.

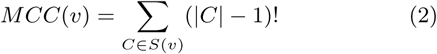

where, *S*(*v*) is the collection of maximal cliques which contain *v*, and (|*C*|− 1)! is the product of all positive integers less than |*C*|. If no edge is present between *v*’s neighbours then MCC(v) is equal to it degree.

## 3. CHEMICAL CHARACTERIZATION OF DNA SEQUENCE: A MAPPING

DNA sequences are combinations of four nucleotides A, T, C, G. Depending upon the chemical structures the four bases can form 16 combinations of dual nucleotides along with repeat bases (AG,GA,CT,TC,AC,CA,GT,TG,AT,TA,GC,CG,AA,TT,CC,GG). Depending upon the chemical structure along with repeating DN groups, the dual nucleotides are classified into seven (07) possible classes [10] given in Table 1. According to the definition of central dogma, transcription and translation are the two steps, through which the information in genes flows into proteins. A protein structure is dependent upon the chemical properties of amino acid by which it is formed. The chemical features of 20 amino acids are basically depending on eight chemical properties, which are shown in Table 2.

**Table 1:** Classification of Dual Nucleotides

| Physicochemical Group | Symbol | Dual Nucleotides |
| --- | --- | --- |
| Purine | R | AG/GA |
| Pyrimidine | Y | CT/TC |
| Amino | M | AC/CA |
| Keto | K | GT/TG |
| Strong H bond | S | GC/CG |
| Weak H bond | W | AT/TA |
| Repeat Group | E | AA/TT/CC/GG |

**Table 2:** Classification of 20 amino acids according to their chemical properties

| Group Name | Amino Acids included |
| --- | --- |
| Acidic | Aspartate, Glutamate |
| Basic | Arginine, Histidine, Lysine |
| Aromatic side chain | Tyrosine, Phenylalanine, Tryptophan |
| Aliphatic side chain | Isoleucine, Leucine, Valine, Alanine, Glycine |
| Cyclic | Proline |
| Sulfur containing | Methionine, Cysteine |
| Hydroxyl containing | Serine, Threonine |
| Acidic amide | Glutamine, Asparagine |

Quantitative understanding of a gene can be made with the help of chemical characterization without any interventions of wet lab experiments. Till date several researchers have tried booms and bursts to make quantitative analysis of a gene family. Some papers are reported where the authors have adopted different graph theoretical approaches to compare several DNA and primary protein sequences [19, 7, 18]. Some authors have tried to make mathematical models on various chemical properties of protein families to understand genes and genome[5, 6].

Analysis of DNA sequences using its underlying biological information hidden within dual nucleotides (DNs) is a good approach and has been reported in some previousre-searches. Randic [13] introduced a qualitative approach to make quantitative comparisons of DNA sequences, whereas, Qi and Fan [11] proposed 3D graphical representation of DNA sequence based on dual nucleotides. They have proposed PN-curve for it. Wu et al. [12] tried to deal with neighbouring nucleotides of DNA sequence.

Our aim is to find out the nature of distribution of physicochemical properties of a DNA sequence in hub genes and its linked genes from the subnetworks of Alzheimer’s. In this regard, it is worth enough to state that, 64 codons which code for 20 amino acids, consists of three nucleotides, where first two of them carry the properties of dual nucleotides.

In this work we try to find out the distribution of physicochemical properties of a DNA sequence from its primary protein sequence in hub and linked genes. Next, we investigate the similarities between linked genes with hub genes which are functionally dependent on hub genes. These investigations may put lights on the basic physical and chemical characteristics of the linked genes of a particular hub genes that may be responsible for Alzheimer’s disease.

We try to investigate physicochemical properties of DNA sequence that keep their signatures in protein synthesis. So, a mapping can be defined between physicochemical properties of any arbitrary DNA sequence and chemical properties of its amino acid sequence. As it is well known that all 16 possible combinations of dual nucleotides built from 4 bases (A,T,C,G) of a DNA sequence exhibits seven (07) different physicochemical properties (R,Y,S,W,A,K,E), which finally converges to the eight (08) chemical properties of amino acid sequences such as Aliphatic, Aromatic, Acidic, Basic, Cyclic, Acid Amide, Hydroxyl, Sulfur, after transcription and translation take place. A possible mapping from DN chemical properties to amino acid chemical properties is shown in Figure 1.

**Figure 1:**
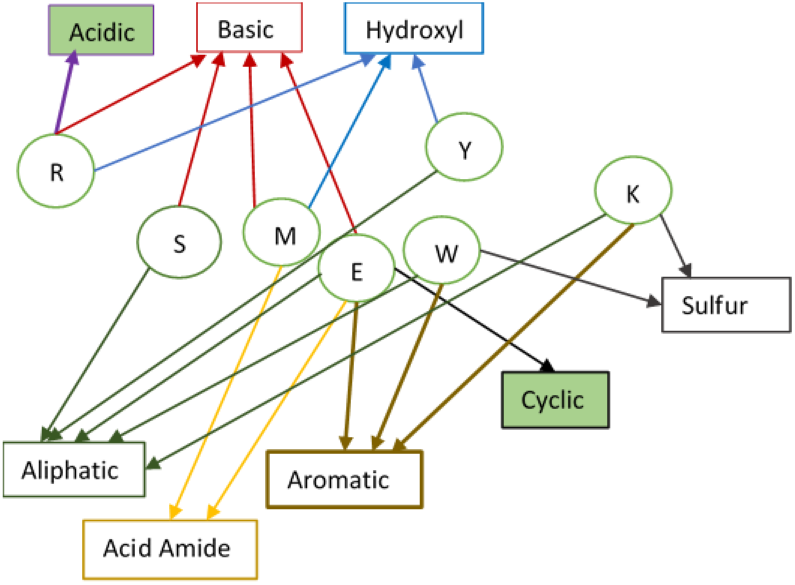
Mapping between physicochemical properties of DNA sequence and chemical properties of amino acid. Dual nucleotide classes are shown in circle and rectangle represents chemical groups

Detail view of the mapping, shown in Figure 2 may give a better understanding of the chemical structures underlying within the three nucleotide positions of the codon.

**Figure 2:**
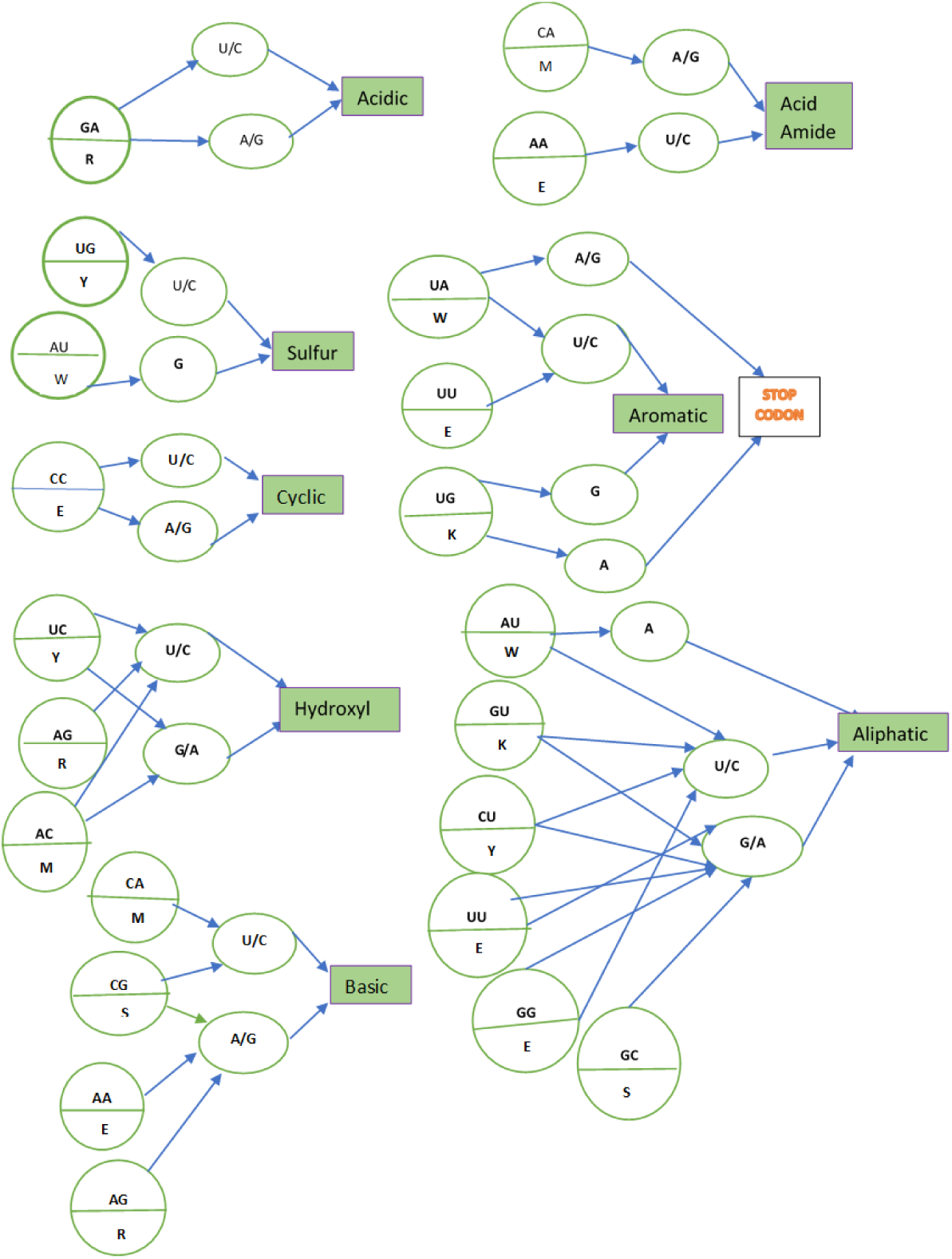
Microscopic view of the mapping between physicochemical properties of DNA sequence and chemical properties of amino acid. The first layer of the mapping represents dual nucleotides, second layer corresponds to nucleotide in third position of the codon and final layer shows the corresponding chemical group (shown inside rectangle) the amino acid belongs to. The corresponding physicochemical groups of DN are shown in half of the circle.

### 3.1 Formal Representation of the Mapping

It is possible to navigate the pathway of chemical properties which is transferred from physicochemical properties of DN sequence to chemical properties of primary protein sequence of itself. Formally, we can define the mapping as follows.

Let 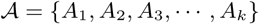 be an arbitrary *k* length nucleotide sequence, where *A_i_* is any nucleotide base i.e. *A_i_* ∈ {*A, T, G, C*}. Assume, 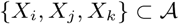 are the triplet nucleotides (codon) derived from given 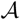, where position of the nucleotides are represented by the lexicographical order of the variables such that *i* < *j* < *k*. Let 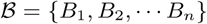 set of seven (07) possible physicochemical properties (R, Y, A, K, S, W, E) of any dual nucleotide. An intermediate mapping function 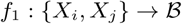 can convert every (*X_i_*, *X_j_*) from any triplet into corresponding physicochemical properties as denoted by 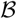.

On completion of this intermediate mapping, while we proceed to traverse to the final mapping, it is important to mention to here that the ordering of (*X_i_,X_j_*) matters biologically. That is, (*X_i_,X_j_*) ≠ (*X_j_, X_i_*) although (*X_i_,X_j_*) and (*X_j_, X_i_*) belong to the group with same physicochemical property from 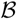. Next, on considering the third nucleotide *X_k_* with the dual nucleotide group, it maps to one of the eight (08)chemical groups of amino acids. Let 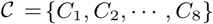 be the set of this eight (08) chemical groups.

Thus, finally we get a mapping from triplet to its corresponding any chemical groups from C by using above intermediate mapping (*f*_1_). We may represent the fact as follows.

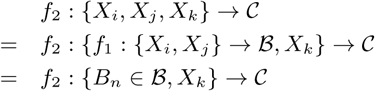

We can now easily map the whole sequence 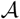 by extracting all sequence triplet {*X_i_,X_j_, X_k_*} and converting it to corresponding to chemical groups using above mapping functions.

### 3.2 Matrix Representation

If we consider the matrix representation of the above fact, then according to Figure 2, it is possible to make 8 × 33 vector representation of each DNA sequence. The vector representation of APP gene is shown in the Table 3, where we can find that Glycine having chemical property of Aliphatic group and coming from repeating group E with relatively high abandance. As it is known that Aliphatic R groups are hydrophobic and nonpolar, those are one of the major driving forces for protein folding, give stability to globular or binding structures of protein [9]. It can be observed that DNA sequence keeps their signature even after translation.

**Table 3:**
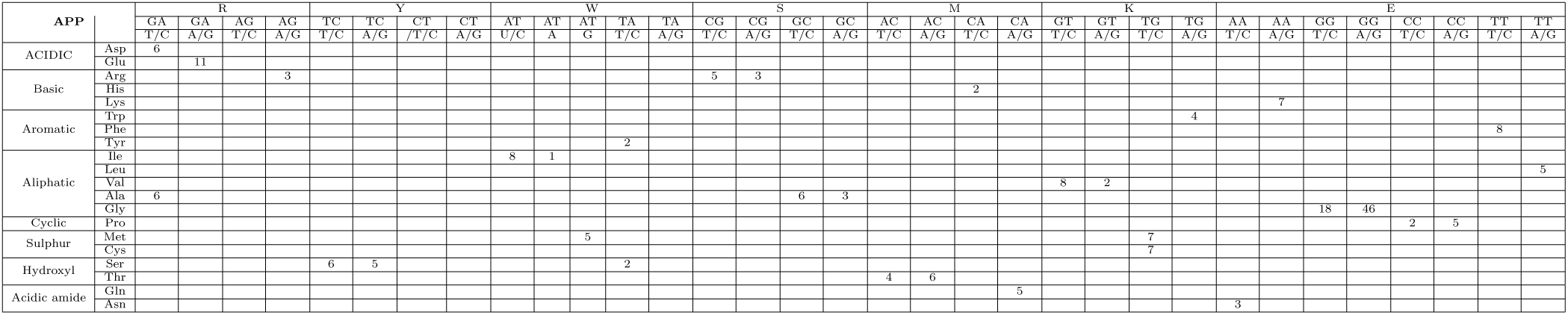
Matrix representation of DNA sequence of APP showing the mapping between physicochemical properties of DN and chemical properties of its primary protein sequence.

## 4. CHEMICAL PROXIMITY BETWEEN SEQUENCES

In this section we try to compute association between DNA sequences of a pair of genes with respect to their distribution of chemical groups mapped from their sequences. The intension behind such proximity computation is to investigate how a hub gene is chemically associated closely with its neighbour or linked genes from its subnetwork.

We read a given DNA sequence in terms of triplets and classify them according to the chemical nature of the amino acid to which it codes. At first, we compute the distribution of chemical groups per sequence from the above matrix representation followed by distance calculation between every neighbour genes with all other hub genes irrespective of any particular subnetwork. We explain the steps below.

### 4.1 Calculating Distribution of Amino Acids

In nature, twenty (20) amino acids are distributed unevenly in a DNA sequence and hence their chemical groups. However, the percentage of abundance of any chemical groups definitely play certain roles in protein structure formation. Hence, it may be responsible for any functional dependencies between a pair of genes. We use the matrix (subsection 3.2) derived after mapping a sequence to corresponding chemical groupings. We calculate the occurrence percentage of each groups with respect to the sequence size. For a given DNA sequence 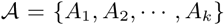, the chemical group distribution can be represented as a vector 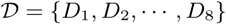 of size eight (08) for eight different chemical groups. Each *D_i_* indicates the percentage of occurrence of a particular chemical group in the given sequence.

### 4.2 Constructing Distance Matrix

Depending upon the distribution of chemical groups of amino acid in hub and linked genes, we calculate a distance between them to see their proximity with respect to their chemical distribution. The functional dependencies between two genes are directly proportional to the degree of similarity of them. Given two chemical group distribution vectors say, 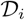 and 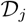, for two target genes, we may calculate the proximity with respect to their chemical properties as follows.

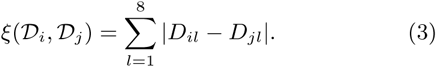

We extend the pairwise distance calculation among all linked genes and all hub genes to get a final distance matrix.

## 5. EXPERIMENTAL RESULTS

We use few selective Alzheimer’s subnetworks and based on their significance in the disease. In this study, six hub genes and their corresponding linked genes which are functionally dependent on them are taken into account to carry out the experiments. The set of hub genes and their neighbours are shown in Table 4.

**Table 4:**
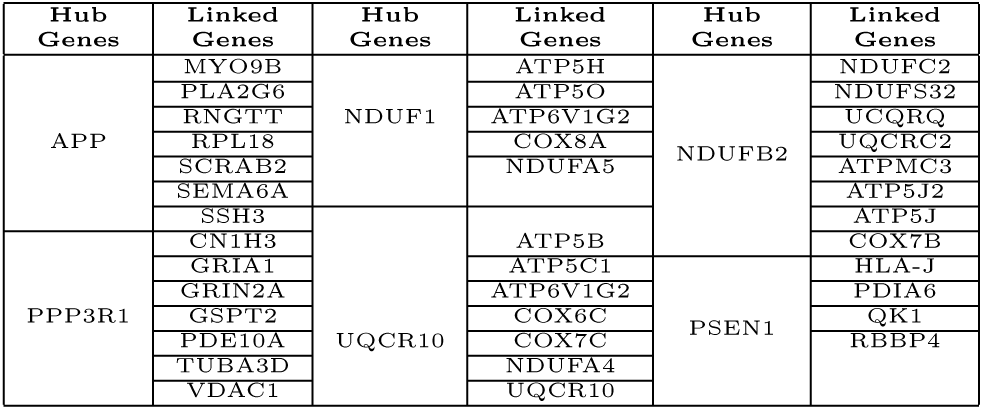
Hub Genes and Linked Genes Table 5: Distance matrix for neighbours of PSEN1 with all other hub genes.

During the analysis of those 6 hub genes and their associated genes it is observed that out of 20 amino acids, amino acids of aliphatic group contribute a large to construct primary protein sequence. Although DNA sequences have six physicochemical properties along with repeating group. It is observed that in most of the cases they come from repeating group (AA/TT/CC/GG) and code for corresponding amino acids. Out of all neighbours of APP, RPL1 shows high enrichment of Glycine which is Aliphatic and comes from GG (repeating group). MYOMB shows high abundance of Proline, which is from cyclic amino acid group but comes from CC (repeating group). APP, the hub gene also has Glycine the most and is from repeating group of physicochemical property and Aliphatic in nature. The chemical distribution of two hub genes PSEN1 and NDUFB2 and their neighbours are shown in the Figures 3 and 4. Interestingly both the figures reveals an interesting pattern of similar chemical group distribution between members of a subnetwork centered around hub genes.

**Figure 3:**
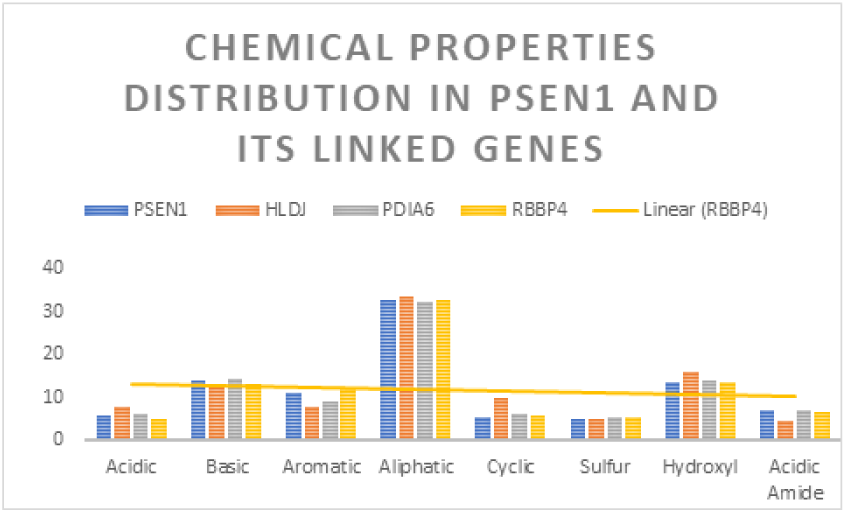
Chemical characteristics of PSEN1 subnetwork

**Figure 4:**
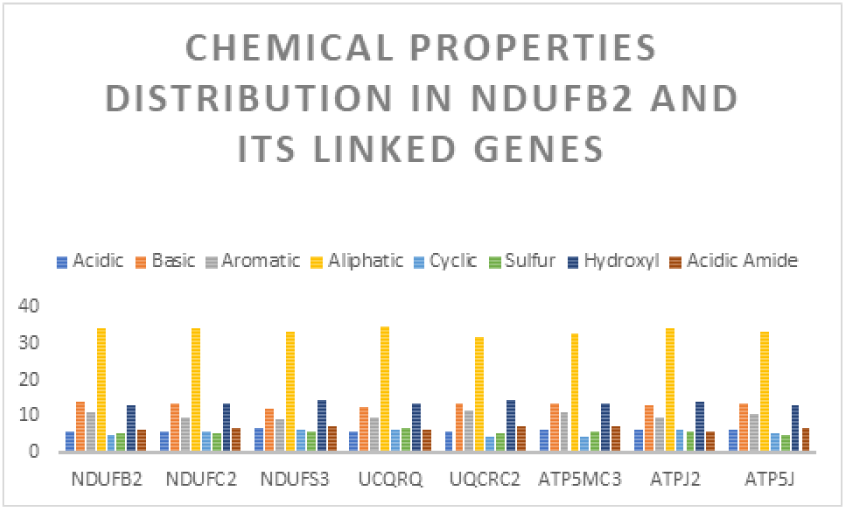
Chemical group distribution of NDUFB2 and its neighbour genes

To investigate the proximity of hub genes with its neighbours we calculate the distance matrix from chemical distributions as discussed in section 4.2. Distance matrix for all the neighbour genes of PSEN1 and NDUFB2 with all other hub genes are shown in Table 5 and 6 respectively. We consider 6 hub genes along with their linked genes to investigate the distribution of chemical properties of amino acid in them and to observe how much similarities do the linked genes have with their hub genes along with their neighbouring hub genes too. We can easily observe from the table that the linked genes of a hub genes PSEN1 and NDUFB2 respectively have smaller distances in comparison to other hub genes. This establish a new and interesting fact that any interacting genes based on their co-expressions are also chemically close to each others. Among 6 networks, hub gene PSEN1 and its three linked genes have strong similarities in distribution of chemical properties of amino acids. But it is also observed that the neighbours of APP such as MYO9B, PLA2G6 and RPL18 showing high proxity with PSEN1 and NDUFB2 instead of APP. So PSEN1 and NBUFB2 may have enough strength to be a hub gene compared to APP for those neighbour genes. It may indicate a fact that association derive from expression data may not always be true from biological point of view.

**Table 5:**
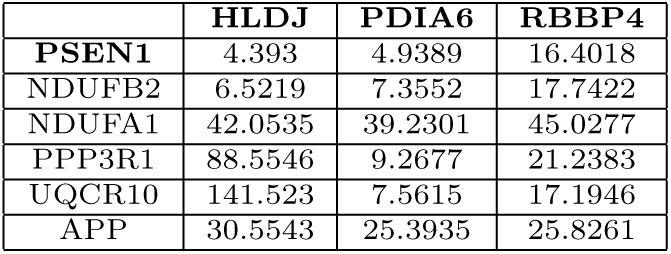
Distance matrix for neighbours of PSEN1 with all other hub genes.

**Table 6:**
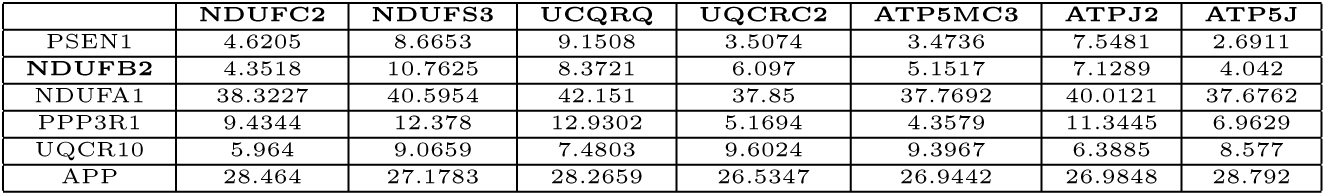
Distance matrix for the neighbours of NDUFB2.

## 6. CONCLUSION

*In silico* inference of gene interaction networks from experimental data is a challenging and important task in computational biology. Majority of the works use expression data as an as a major source of input to infer the network. We believe that during interaction the chemical properties of the amino acids and their distributions in a gene plays a vital role. In this work we demonstrated the justification of the fact by applying our idea in a selective subnetworks of Alzheimer’s disease networks which is derived from gene expression data. We observed that chemical properties are distributed very similar way between a hub gene and its neighbours. We proposed a new mapping technique for converting a DNA sequence to corresponding amino acids chemical groups. It is our firm belief that our investigation will give a new dimension in the future research on computational network inference.

## Footnotes

1 https://www.ncbi.nlm.nih.gov/geo/

